# Insect behavioral evidence of spatial memories during environmental reconfiguration

**DOI:** 10.1101/118687

**Authors:** Diogo Santos-Pata, Alex Escuredo, Zenon Mathews, Paul F.M.J. Verschure

## Abstract

Insects are great explorers, able to navigate through long-distance trajectories and successfully find their way back. Their navigational routes cross dynamic environments suggesting adaptation to novel configurations. Arthropods and vertebrates share neural organizational principles and it has been shown that rodents modulate their neural spatial representation accordingly with environmental changes. However, it is unclear whether insects reflexively adapt to environmental changes or retain memory traces of previously explored situations. We sought to disambiguate between insect behavior at environmental novel situations and reconfiguration conditions. An immersive mixed-reality multi-sensory setup was built to replicate multi-sensory cues. We have designed an experimental setup where female crickets Gryllus Bimaculatus were trained to move towards paired auditory and visual cues during primarily phonotactic driven behavior. We hypothesized that insects were capable of identifying sensory modifications in known environments. Our results show that, regardless of the animals history, novel situation conditions did not compromise the animals performance and navigational directionality towards a novel target location. However, in trials where visual and auditory stimuli were spatially decoupled, the animals heading variability towards a previously known location significantly increased. Our findings showed that crickets are able to behaviorally manifest environmental reconfiguration, suggesting the encoding for spatial representation.

## Introduction

Insects robustly navigate within very dynamic environments with limited resources. At the sensorimotor level, insects need to integrate multi-modal stimuli and constantly adapt their behavioral strategies in order to survive. Their relatively simple nervous system has been subject of research aimed at revealing the mechanisms behind insect navigation and often their distributed nervous system is described as a stack of simple reflexive loops.

Goal-oriented behavior has been extensively observed in insects such as chemotaxis performed by the silkmot^1,2^ or phonotaxis by female crickets^3^. Earlier research suggests that parallel direct sensory-motor loops are supplemented by specific brain areas that serve to integrate functions related to context, learning and smooth coordination of action^4^. However, if insects would simply perform reactive behaviors, they could not be able to recall hidden target positions^4,5^, or to form sensory-action couplets associations^5,6^. Kraft and colleagues [2011] have shown that flying bees are capable of using polarized-light information to direct their routes towards a food location. Also, experimental work has been conducted to explore the capabilities of flying insects to navigate within distinct maze configurations. In^7^, the authors have found that the performance of bees navigating within a maze depends on the regularity of the maze, suggesting a process of spatial abstraction rather than memorizing entire navigational sequences. Such form of context-dependent learning leads to the question of whether insect behavior, as for mammalian behavior, could be described through the main components of operant conditioning paradigms^8^.

Operant conditioning implies the ability to build relational maps of contexts, actions and expected outcomes in order to boost decision-making processes. This form of learning can be observed in insect behavior such as honey-bees representing food locations (waggle dance)^9^ or communication suppressing through stop signals by partners who have experienced conspecific attacks at a food source^10^. Furthermore, it has been shown that insects are capable of performing logical operations such as numerical counting^11^, pattern recognition^12^ or context dependent learning^13^ and most of their behaviors imply successful integration of different sensory modalities signals. These results advocate that insects might be capable of developing higher level representations of not only their surroundings but also other agents sharing their territories.

In the domain of spatial representation, insects make use of allocentric environmental information to perform homing tasks and it has been suggested that visual landmarks are used to form cricket place memories^4^. In this context, emphasis is often placed on the role of the mushroom bodies in the integration of sensory and motor signals. Indeed, neural responses in the mushroom body appear to be related to specific directions of the animal’s orientation, which constitutes a type of spatial representation of their body in space^14^.

Despite is growing evidence for insect environmental adaptation and place-memory formation during navigational tasks, it still remains unclear whether insects form memories of context specific sensory statistics or simply perform stimulus-stimulus associations. On one hand, behavioral adaptation could be a reflex derived from environmental reconfiguration. On the other hand, behavioral modulation could be explained by higher level functions such as perceiving contextual modifications.

We sought to find out the behavioral specifics of insects exposed to either environmental novel situations or environmental reconfiguration. Novel situations are characterized by changes in the animals spatial target location while environmental reconfiguration maintain the animals goal location but with induced sensory modifications. We hypothesized that insects are able to form representations of environmental sensory statistics and display behavioral signatures of environmental sensory reconfiguration. Crickets have a specialized nervous system and behavior repertoire dedicated to sound communication^15^, making them ideal experimental subjects to test modulation of goal-oriented behavior. Male crickets periodically move their wings, rubbing a flexible membrane at the tip of one wing called *plectrum* against a set of *files* placed under the opposite wing^16^, generating an attractive species-specific mating sound. Females, on the other hand, possess tympanal membranes in their forelegs allowing them to orient towards the bursting sound of a potential mate, a type of behavior called phonotaxis behavior in which acoustic information is the main component of goal-directed navigation.

In order to access memory traces of reconfiguration, we tested female crickets during stimulus-stimulus associations tasks and we have quantified their adaptive stereotypical behavior after stimulus-stimulus dissociation while performing goal-oriented navigation.

## Methods

### Experimental set-up to study insect navigational behavior

Besides experimental research to study insect behavior within physical arenas^17–19^, a number of different paradigms have been proposed to analyze animal behavior. On one hand, locomotion compensatory apparatus have been extensively used to study insect navigation since the 1970’s^19–21^. Also, open-loop systems were proposed to analyze cricket antennae movements during visually guided behavior^22,23^. Similar approaches have also been generalized to study neural mechanisms involved in rodents spatial representation^24^. Insect hybrid systems have been deployed to study insect behavior in a range of navigational tasks such as phonotaxis behavior^25,26^. Having been tested in a myriad of experimental studies, locomotion compensatory apparatus appear to be a plausible tool for investigating animal navigation within controlled environments. In combination with mixed-reality environments, these technologies increase the amount, complexity and ecological validity of possible stimuli to be used during experiments. To study animal behavioral correlates of spatial memories in novel situations and reconfiguration conditions we have built a virtual reality immersible system (Figure 1-B). The apparatus consists of a polystyrene ball with a diameter of 8 cm floating on an air cushion allowing a tethered insect to move on top of it. The insect is tethered to a rigid beam in such a way that it remains placed on the floating sphere with fixed position and orientation, but perceiving coherent visuo-motor responses from the virtual environment. Two two-dimensional optical sensors, located on the horizontal plane and perpendicular to each other, were used to track the sphere’s yaw, pitch, and roll transformations triggered by the insect’s gait. Further, captured animal movements were mapped to navigational displacements within the virtual environment. Three computer screens surrounded the sphere providing 270^*∘*^ visual stimuli to the animals. Computer screens were Samsung SyncMaster 943NW (LCD), 1440×900 pixels in resolution, 17.3 inches width and 14.5 inches height, with a refresh rate of 75Hz and 300cd/m2 of image brightness. A virtual environment was developed using the 3D Object-Oriented Graphics Rendering Engine (OGRE), where landmarks, ambient light or other graphical properties were programmatically manipulated. A sound application (PureData) was integrated into the system to generate insect-specific mating-sounds. Two micro-speakers were placed at a distance of 35cm from the animal and at ±45^*∘*^ azimuth to the front-screen display, providing stereo sound stimuli to the animal tympanal membrane^27^. Visual and auditory sensory modalities together with motor actions were synchronized within the virtual reality environment.

**Figure 1.**
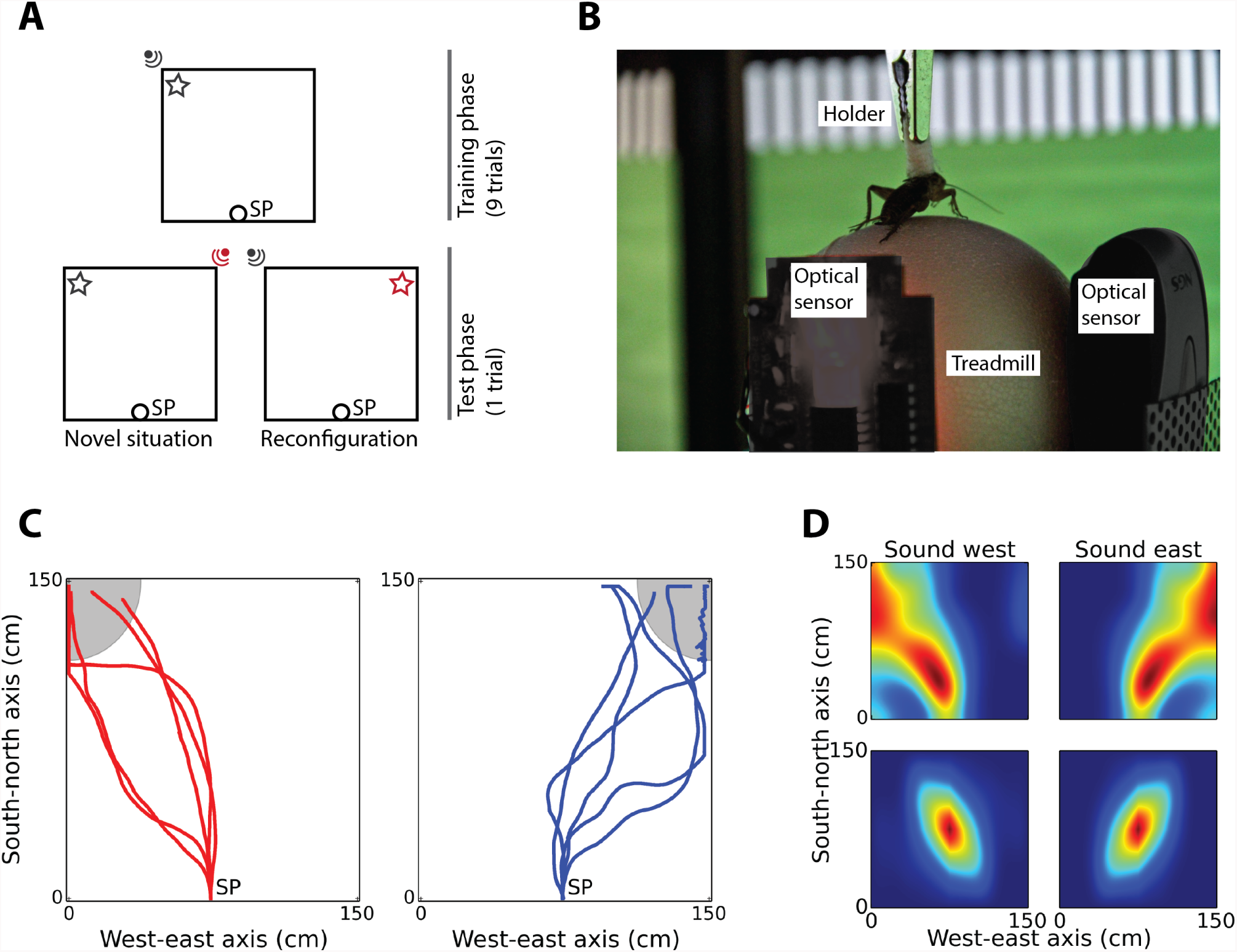
Experimental protocol and setup. **A** Illustration of the experimental protocol. Trajectories begun at starting position (SP) until animals reached the target location (sound). Training trials (top) consisted of paired locations of visual and auditory stimuli. At the testing trial (bottom), either the auditory (novel situation) or the visual stimuli (environmental reconfiguration) was moved to a novel position contra-lateral to its testing position. **B** Snapshot of the treadmill setup surrounded by the computer displays. A female cricket runs on a floating sphere’s treadmill sustained by a holder. Sphere rotations are captured by two optical sensors and translated into the virtual reality engine, providing coherent sensory-motor feedback. **C** Phonotaxis behavior with sound source at north-west and north-east locations of the virtual environment. **D** Group rate-maps after Gaussian kernel interpolation with 10 pixels standard deviation (top) and occupancy autocorrelograms (bottom) for sound at north-west and north-east conditions.

### Experimental design

For the experiments, we acquired female crickets *Gryllus Bimaculatus* with undetermined age bought in colony-boxes from a local store. Animals were fed daily with fruit-based diet, maintained at room temperature and an effort was made to keep the light/dark regime fixed at 12:12-h LD cycle. Animals were food deprived and kept in separated transparent Perspex boxes (20 × 35 × 20 cm) 24 hours before the experiment. The animals were trained with sound and visual stimuli spatially paired, with each trial consisting of navigating from a starting position at the south-center location to the north-west location of the virtual environment (9 trials per animal, figure 1-A top) and was deemed completed when animals reached at least 80 percent of the south-north axis length (1.5 * 0.8 meters). In order to induce novel situation or reconfiguration, at the tenth trial one of the stimulus was moved to a novel position (north-east location of the virtual arena). Because sound is the main drive of action during phonotactic behavior^3^, placing the sound source at a different position (north-east), which implies changing of the goal-location, was considered as the novel situation condition (Figure 1-A, bottom-left). On the other hand, because the visual cue was the stimulus associated to sound but did not affect the target location, moving the visual landmark to a novel position (north-east) was considered as environmental reconfiguration (Figure 1-A, bottom-right). With the current setup, we have trained the animals (9 trials each) to move from the starting position towards the north-west location where sound and visual cues were paired (figure 1-A). At the tenth trial, we have replaced either the sound or the visual cue at the north-east corner of the virtual arena. Animals (n=10) experiencing a displacement of the sound-source to a novel location during the testing trial belonged to the novel situation group, while animals (n=10) experiencing a displacement of the visual cue to a novel location at the testing trial were considered as belonging to the environmental reconfiguration group. In between trials, animals were allowed to rest for approximately 30 seconds and rewarded with fruit-sucrose, independently of their navigational performance.

In our setup, in order to motivate female crickets to navigate towards a species-specific mating sound, during each trial, a 4.8 kHz sound tone at a syllable rate of 30 Hz was generated in real-time to mimic male cricket species-specific calling song^16^. The sound amplitude of each loudspeaker was determined by the Euclidean distance and heading orientation between the animal location and the sound-source location within the virtual environment arena. After the experiment, the animals were released in the open field surrounding the university campus.

### Navigational quantification

Measures of navigational performance focused on both trajectory directionality and angular variability. Trajectory directionality provides a quantification of the expressed goal-directed behavior that insects had to perform during the experiments and was accessed by extracting the angular components of individual trajectory segments. Angular variability was computed through the variance of segments orientation. We have also extracted occupancy measures for group trajectories, which provides a quantification for direction coherence among individual belonging to the same group (figure 2-C). In order to quantify for the overall group occupancy metrics, we have extracted the first spatial contour above 10 percent of the occupancy correlograms maximum activity and calculated their eccentricity. To do so, we computed the semi-major vector between pairs of points found in the occupancy contour as:

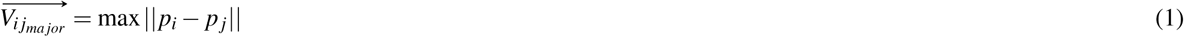

where *ij* represent every pair of points along the autocorrelogram contour and || the euclidean distance between *i* and *j*. After finding the contour longest vector 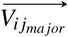, we applied a projection from its center point with an angular orientation of 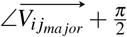 obtaining a projection 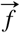 used to define 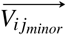 through its intersection with the ratemap occupancy contour, such that:

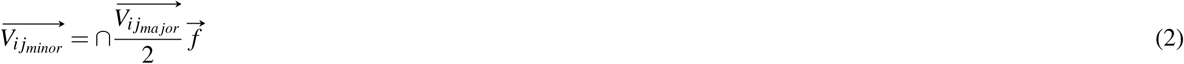

**Figure 2.**
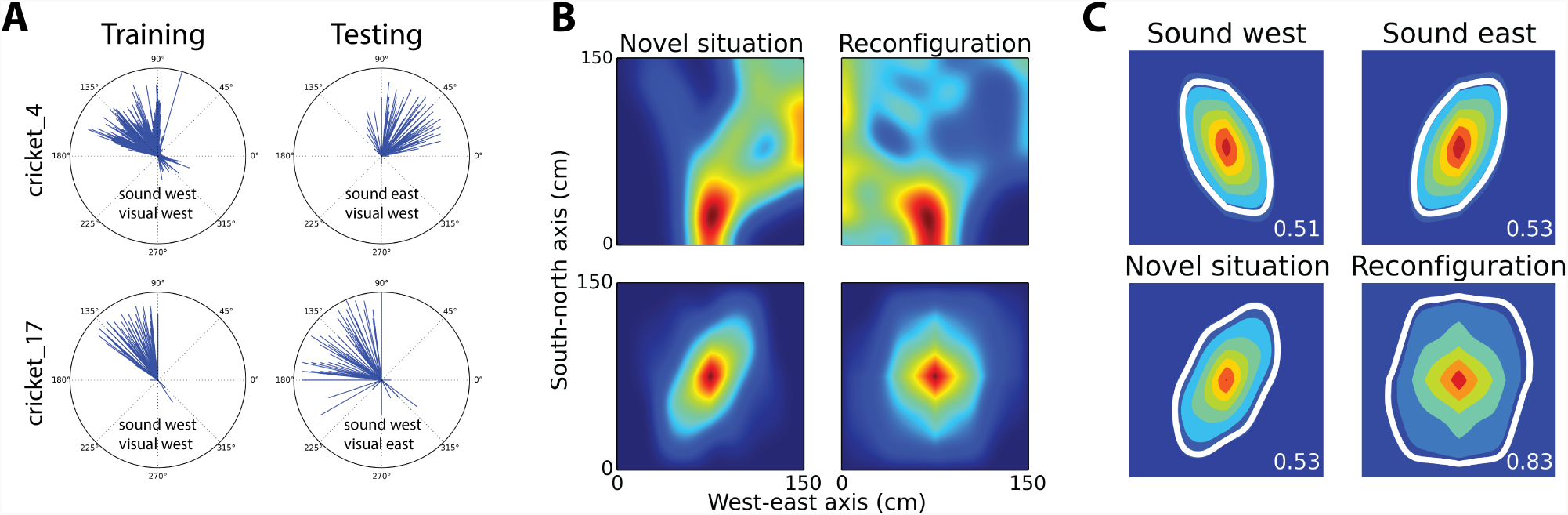
Experimental protocol and setup. **A** During training (left) both groups exhibited directed movements towards the north-west location of the virtual environment. At testing trials (right), the novel situation group (top-right) shifted their directionality towards the north-east location. **B** Group rate-maps after Gaussian kernel interpolation with 10 pixels standard deviation (top) and occupancy maps (bottom) at novel situation and environmental reconfiguration conditions. **C** Occupancy maps eccentricity measurement. Environmental reconfiguration condition scored lower for directional specific navigation.

The occupancy autocorrelogram eccentricity (OAE) which provides a measure of movement directionality was given by:

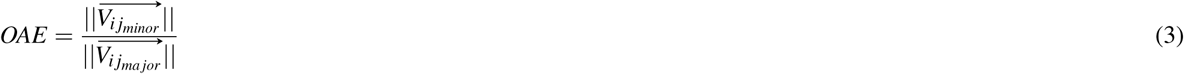

allowing the extraction of the overall group directionality during phonotaxis navigation.

## Results

Prior to the experiments and in order to validate induced visual stimuli we quantified the antennae reaction response of 10 female crickets to visual cues traversing within the virtual environment. During testing, animals remained steady on top of the spherical treadmill; however, their spontaneous motor responses were not translated into motor responses within the virtual environment. Visual cues were presented within ±120 angular degrees of the animal head-orientation and randomly interleaved between the three computer screens surrounding the animal. Antennae responses to visual cues were visually assessed and marked as valid when at least one antennae coherently followed the trajectory of the visual cue. Overall, visual cues appearing in the centered computer screen triggered higher responses (95%±4%, mean±std) than lateral computer screens (79%±9%, mean±std). Nonetheless, it suggests that the crickets were able to perceive visual cues within the virtual environment.

In order to validate the effectiveness of the sound stimulation on guiding the animals behavior, we conducted a between-groups analysis of their navigational trajectories final locations. Trajectories of both groups (n=5 per group) started at the south-center location (SP, figure 1-A) and, as for the main experiment, were considered as valid when animals reached at least 80 percent of the south-north axis length. For the first group, the sound-source was placed at the north-west location of the virtual environment, while for the second group the sound-source was placed at the north-east location. Results revealed a significant difference in position along the horizontal axis between the two groups (Wilcoxon rank-sum test: *p =* 0.02), which suggests that the synthesis and mapping of the auditory stimuli was sufficient to modulate animal goal-directed behavior (figure 1-c, 1-D). Coherently, overall group navigational ratemaps and occupancy autocorrelograms suggested effectiveness in modulating the animals goal-location and trajectory directions (figure 1-D). Given the induced behavioral modulation through the sound stimuli, we next tested cricket navigational behavior in both novel situation and reconfiguration conditions.

In order to quantify navigational directionality, we have extracted the overall occupancy maps from group trajectories belonging to each condition (figure 2-C). Visual inspection of occupancy ratemaps suggested that animals successfully navigated towards the sound source location. However, despite the fast adaptation to a novel target location during the novel situation condition, the navigational occupancy for environmental reconfiguration condition revealed a higher spreading. Indeed, ratemaps autocorrelograms showed differences in their occupancy resolution between novel situation and reconfiguration conditions (figure 2-C).

Quantification of navigational directionality revealed a lower OAE score for sound-only navigation (north-west *OAE* = 0.51; north-east *OAE* = 0.53) and novel situation group (*OAE* = 0.53) when compared with the environmental reconfiguration group (*OAE* = 0.83) (figure 2-C). Higher occupancy autocorrelogram eccentricity depicted lower navigation directionality for the reconfiguration group, suggesting a loss in the ratio of directed movements towards the goal-location.

In addition to navigational occupancy and eccentricities measures, we analyzed changes in navigational directionality performed across groups. Because within the presented setup trials could be successfully accomplished through vector-based navigational strategies, we extracted trajectory segments from the last training and testing trials for both groups. Segments extraction was performed through a sliding window with bin size of 10 data points in the trajectory array and by computing the vector between first and last position within each bin. Segments with magnitude zero, when animals were quiet, were removed from the analysis. Examples of head orientation of two animals belonging to different groups can be appreciated in figure 2-A. For novel situation conditions, the animals heading movements have shifted to the contra-lateral angular quadrant during the testing trial (e.g. cricket 4), while for environmental reconfiguration the animals heading movements were maintained at the testing trial (e.g. cricket 17), again revealing the effectiveness of the sound stimulation during phonotactic behavior.

To quantify changes in navigational directionality between the last training and testing trials in both groups, we have compared the head-orientation distribution along angular coordinates (figure 3-A). In accordance with the experimental design, the distribution of segments orientation were more prominent between 54 and 126 degrees, which represents the angular interval to perform vector-based navigation towards a target position placed either at the north-west or north-east corners of the virtual arena. Differences in orientation distribution between last training and testing trials revealed higher changes for the novel situation group than for the reconfiguration group. That was justified by the fact that the sound-source at the testing trial was moved from north-west to the north-east location for novel situation but not for the reconfiguration group.

**Figure 3.**
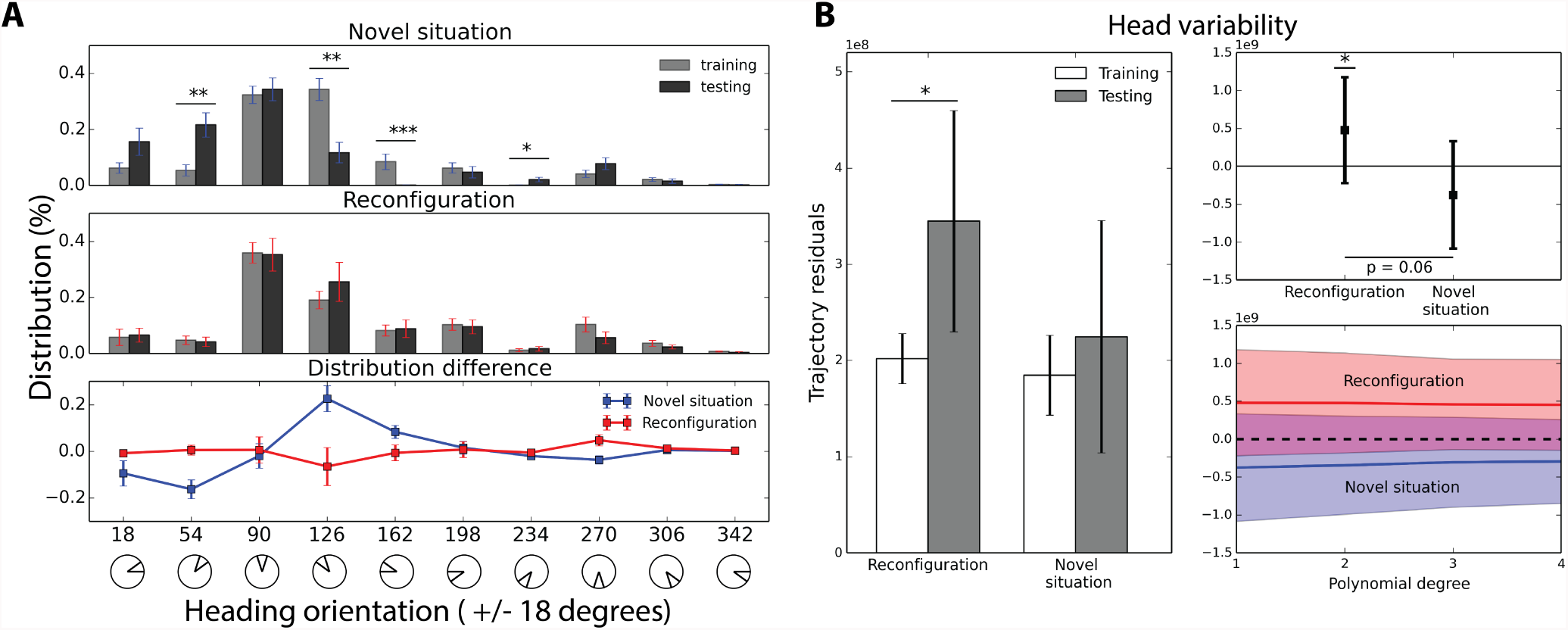
Experimental protocol and setup. **A** Movement orientation distribution. Comparison between last training and testing trials revealed significant differences for the novel situation group (top panel), but not for the environmental reconfiguration group (middle panel). Each segment encodes head orientation range of 36°. Angular distribution differences between last training and testing trial for both groups (bottom panel) **B** Trajectory variance. Variability of the trajectory obtained from the sum of squared residuals of the least squares linear fit rank (left and top-right panels). Comparison of residuals between last training and testing trials revealed to be significant for the environmental reconfiguration group (t-test *p* = 0.051). Trajectory residuals was independent of the fit polynomial degree (bottom-right panel), suggesting that variability was not an artifact of vector/non-vector-based navigation type.

As expected, the group experiencing novel situation during the testing trial, where the sound-source was moved to a novel location, exhibited greater differences for segments distribution at 54 and 126 degrees. Specifically, animals showed preference for orientations towards 126 degrees during training (Mann-Whitney U test *p* < 0.01), while during testing animals head orientations aligned towards 54 degrees (Mann-Whitney U test*p* < 0.01). On the other hand, movement heading orientation distribution showed no significant differences between training and testing for the environmental reconfiguration condition, where the sound-source but not the visual-stimulus remained at the same location.

We have observed that the occupancy autocorrelogram eccentricity was higher for reconfiguration than for novel situation conditions (figure 2-C). Because head orientation differences *per se* are not sufficient to determine if there is lack of perceptual information regarding environmental reconfiguration, we have assessed the animals navigational variability from their trajectories.

To quantify group navigational variability, we performed a linear least-squares fit on the animals trajectories at both last training and testing trials. Within our setup, optimal performance could be achieved through vector-based trajectories. Thus, the linear fit residuals provided a measurement for trajectory variability (figure 3-B). As anticipated by the occupancy autocorrelogram eccentricity measures, the trajectory variability at testing trials was significantly different for environmental reconfiguration but not for the novel situation condition (t-test *p* = 0.051), suggesting that movement intentionality was lower when the sensory configuration was altered (figure 3-B). Normalization to baseline (subtraction of residuals at training trials with the ones at testing trials) revealed a partially significant difference between groups (Ranksum test *p* = 0.06, figure 3-B top-right panel). Because differences in navigational variability could be an artifact of a linear-fit onto a non-vector-based navigational performance, we have quantified group differences up to 4 polynomial degrees which would reflect arc trajectories. However, group differences were independent of the polynomial degree, suggesting that navigational variability was independent on the type of trajectory performed (figure 6 bottom-right).

## Discussion

Insects display striking skills for spatial exploration and are capable of effortless adaptation to dynamic environments. Numerous studies have focused on insect place-memory formation revealing their ability to integrate environmental information. Wessnitzer and colleagues (2008) have shown how crickets are able to improve their navigational performance in an aversive dry version of the Morris-water maze^28^ by considering visual cues to situate themselves in space. Moreover, in the same experimental setup, it was suggested that crickets were able to locate a remapped hidden spot by following previously learned spatial-sensory cues contingencies. Because insects quickly adapt to dynamic environments, one could argue that their reflexive capabilities would be sufficient for surviving in such territories. Another possibility would be that insects are capable of recognizing the environment’s dynamics instead of considering each experience as occurring within an unique spatial context. Thus, there is no sufficient evidence of whether insect spatial adaptation is a mnemonic property or a consequence of environmental modifications.

It has been previously shown that rodents modulate the rate of hippocampal neural activity during environmental morphing^29^, suggesting that mice constantly update their internal spatial representation^30,31^. The neural structures of mammals and insects have been compared and suggested to share common organizational principles^32,33^. Also, it has been suggested that the genetic mechanisms involved in pattern formation of the embryonic body plan and brain development are shared for both insects and mammals^34^. Thus, one could expect that the neural mechanisms underlying insect navigation and spatial representation would resemble the ones of rodents.

Here, we have made an effort to understand whether insects, like rodents, are able to contextualize themselves in space and integrate environmental modifications into their behavioral plans. Within our experimental design we have focused on insect navigational adaptation during novel situation and environmental reconfiguration scenarios during goal-oriented behavior. To do so, we have trained female crickets *Gryllus Bimaculatus* to perform phonotaxis behavior within a virtual environment and navigate towards a target location where a visual cue was spatially paired with the sound stimulus. After training trials crickets experiencing the novel situation condition quickly adapted their navigational plans by reorienting their learned navigational movements towards a novel target location. Contrarily, crickets experiencing the environmental reconfiguration condition have maintained their target-location (figure 3-A). However, their movements orientation variability towards the target location significantly increased when compared to variability of the novel situation group. Our results suggest that crickets were able to maintain an internal representation of their surroundings and behaviorally expressed environmental modifications through their movements orientation variability (figure 3-B).

Internal representations of space are supported by multiple neural mechanisms in the rodent brain, such as place-cells, grid-cells and head-direction-cells^30,31,35,36^. Similarly, head-direction-cells have been found in the insect mushroom bodies^37^. However, it is known that insects do not perform only vector-based navigation, raising the possibility that they might also use similar neural mechanisms for spatial representation as rodents do (see^38^ for a review).

Insects and rodents perform a common panoply of behaviors, such as homing, foraging and hoarding. Indeed, honey bees perform magnificent hoarding tasks by navigating towards abundant pollen locations and bringing the nectar back to their hives^39,40^. Similarly, desert-hamsters exploit their navigational apparatus to successfully store resources at their home location. However, while the neural circuitry involved in mammalian navigation is relatively well understood and further supported by computational models implementing the neural components necessary for hoarding behavior^41^, the computational processes involved in insect navigation is far-less understood. Therefore, how much of those mammalian neural components can be validated into a biologically plausible insect cognitive-architecture, still remains an open question. Moreover, cognitive processes thought to be found only in higher order animals, might also have developed in simpler nervous systems animals, such as insects.

## Acknowledgements

The research leading to these results has received funding from the European Research Council under the European Union’s Seventh Framework Programme (FP7/2007–2013)/ERC grant agreement no [341196] cDAC.

## Author contributions statement

Z.M. and P.V. conceived the experiment, D.S.P. conducted the experiment and analyzed the results. D.S.P, Z.M. and A.E developed the setup. All authors were involved in revision of the manuscript and discussion.

